# Hyphal aggregation and surface attachment of *Streptomyces* is governed by extracellular poly-β-1,6-*N*-acetylglucosamine

**DOI:** 10.1101/089920

**Authors:** Dino van Dissel, Joost Willemse, Boris Zacchetti, Dennis Claessen, Gerald B. Pier, Gilles P. van Wezel

## Abstract

Streptomycetes are multicellular filamentous microorganisms, which are major producers of antibiotics, anticancer drugs and industrial enzymes. When grown in submerged cultures, the preferred enzyme producer, *Streptomyces lividans*, forms dense mycelial aggregates or pellets, which requires the activity of the proteins encoded by the *matAB* and *cslA-glxA*. Here we show that matAB encodes the biosynthetic genes for the extracellular polymeric substance (EPS) poly-β-1,6-N-acetylglucosamine or PNAG. Heterologous expression of matAB in actinomycetes that naturally lack these genes was sufficient for PNAG production and induction of mycelial aggregation. Also, overexpression of *matAB* in a non-pelleting *cslA* mutant restored pellet formation, which could effectively be antagonized by the PNAG-specific hydrolase, dispersin B. Extracellular accumulation of PNAG allowed *Streptomyces* to attach to hydrophilic surfaces, unlike attachment to hydrophobic surfaces, which involves a cellulase-degradable EPS produced by CslA. Altogether, our data support a model in which pellet formation depends on hydrophilic interactions mediated by PNAG and hydrophobic interactions involving the EPS produced by CslA. These new insights may be harnessed to improve growth and industrial exploitation of these highly versatile natural product and enzyme producers.

## INTRODUCTION

The ability of many microorganisms to organize themselves into biofilms has a huge impact on human society, impacting human health (1), waste treatment (2) and crop production (3). Within a biofilm many different individual cells aggregate into a multicellular community where they coexist in a relatively complex and coordinated manner (4, 5). The cells are held together by an extracellular matrix, often referred to as extracellular polymeric substance (EPS), that can comprise to 90% of the mass in a biofilm (6, 7).

The presence of extracellular polysaccharides in the matrix is in most cases essential for a persisting biofilm. Although many different kinds of exo-polysaccharides are employed by different bacterial species (8, 9), pathogenic bacteria often produce poly-β-1,6-*N-* acetylglucosamine (PNAG) to stick to the biotic surface of a host, which often is a requirement, but not by itself sufficient for biofilm formation (Wang *et al.*, 2004, Roux *et al.*, 2015, Mack *et al.*, 1996, Beenken *et al.*, 2004). Interestingly, the soil bacterium *Bacillus subtilis* also produces PNAG, suggesting that this EPS is abundantly present in natural matrices (10). In *Staphylococcus epidermis,* the organism in which PNAG was first detected, the *icaADBC* gene cluster encodes proteins responsible for the production of PNAG. IcaAD form a glycosyltransferase that synthesizes the PNAG chain intracellularly, while IcaB partially deacetylates the polymer extracellularly, thereby changing the net charge, and allowing better association with the cell surface (11). IcaC likely plays a role in the export and possibly O-succinylation of the PNAG polymer (12).

Streptomycetes are multicellular filamentous bacteria that reproduce by sporulation and are the source of the majority of the antibiotics as well as many other compounds of medical, agricultural and biotechnological importance (13, 14). The production of such natural products is under extensive genetic and morphological control (15, 16). In submerged cultures, many *Streptomyces* species form pellets, which may be regarded as self-immobilizing biofilms (17). Because of their dense architecture, which among other issues, causes significant mass transfer limitations, pellets are often undesirable for industrial production; however, antibiotic production often benefits from mycelial clumps compared with mycelial fragments (18, 19). Pellet architecture depends on genetic and environmental factors, like the septum formation (20, 21), cytoskeleton (22), shear stress (23), pH (24) and cell wall fusions (25). Two extracellular polysaccharides are of particular important for cellular aggregation, namely a cellulose-based EPS that is produced by the concerted action of CslA, GlxA and DtpA (26-29), which mediates attachment in cooperation with the amyloid-forming chaplin proteins, and a putative second extracellular polysaccharide that is synthesized by the MatAB proteins (30). Loss of either system results in similar dispersed morphology, suggesting that both systems might work in unison in a so far unclear way (31).

The *matAB* gene cluster shows significant resemblance to the PNAG biosynthetic gene cluster *icaADBC* from *S. epidermidis*, with the bifunctional MatB likely corresponding functionally to both IcaA and IcaB, forming an intracellular glycosyltransferase domain and an extracellular oligo-deacetylase domain, respectively. SCO2961, which is located directly downstream of *matB*, is an orthologue of *icaC* that might also play a role in formation of the mature polymer. The function of MatA is unclear as it lacks known functional domains, but as deletion of the *matA* gene reduces hyphal aggregation it may assist in efficient polymerization of the EPS, similarly to IcaD (32).

In this study we show that MatAB is responsible for the production of PNAG. The MatAB-dependent EPS is required for adherence to hydrophilic surfaces, while a second EPS produced by the action of CslA and GlxA mediates attachment to hydrophobic surfaces. The combination of these two systems make up the architecture of a pellet, creating a strong, robust structure. Since natural product formation and enzyme production depend strongly on the morphology of the mycelia, this also has major implications for biotechnological exploitation.

## RESULTS

### Mat facilitates the formation of a granular layer on the outside of the hyphae

Previous studies showed that the *mat* genes encode a putative extracellular polysaccharide synthetase system in *Streptomyces* that is required for mycelial aggregation and pellet formation in submerged cultures, most likely by mediating cell-cell bonding (30). To elucidate the underlying mechanism, we here investigated the cell surface mechanism by which this is mediated, aiming t elucidating the nature of the EPS. Although the Mat proteins are expressed throughout growth (31), the Mat polymer was most apparent in young mycelia. High resolution imaging by cryo-scanning electron microscopy (SEM) revealed a surface layer that decorated the entire outer surface of the hyphae (Figure 1A). Transmission electron microscopy (TEM) of Tungsten acid-negative stained cells, which images electron dense polymeric surface structures, highlighted an extracellular surface layer (Figure 1E). Between the hyphae a deposit of extracellular matrix could also be observed by SEM (Figure 1C). Conversely, the hyphae of the *matB* mutant have instead a smooth surface, observed both with SEM (Figure 1B) or negative staining in the TEM (Figure 1F). We also failed to detect any extracellular material between the hyphae of *matB* mutants (Figure 1D).

**Figure 1.**
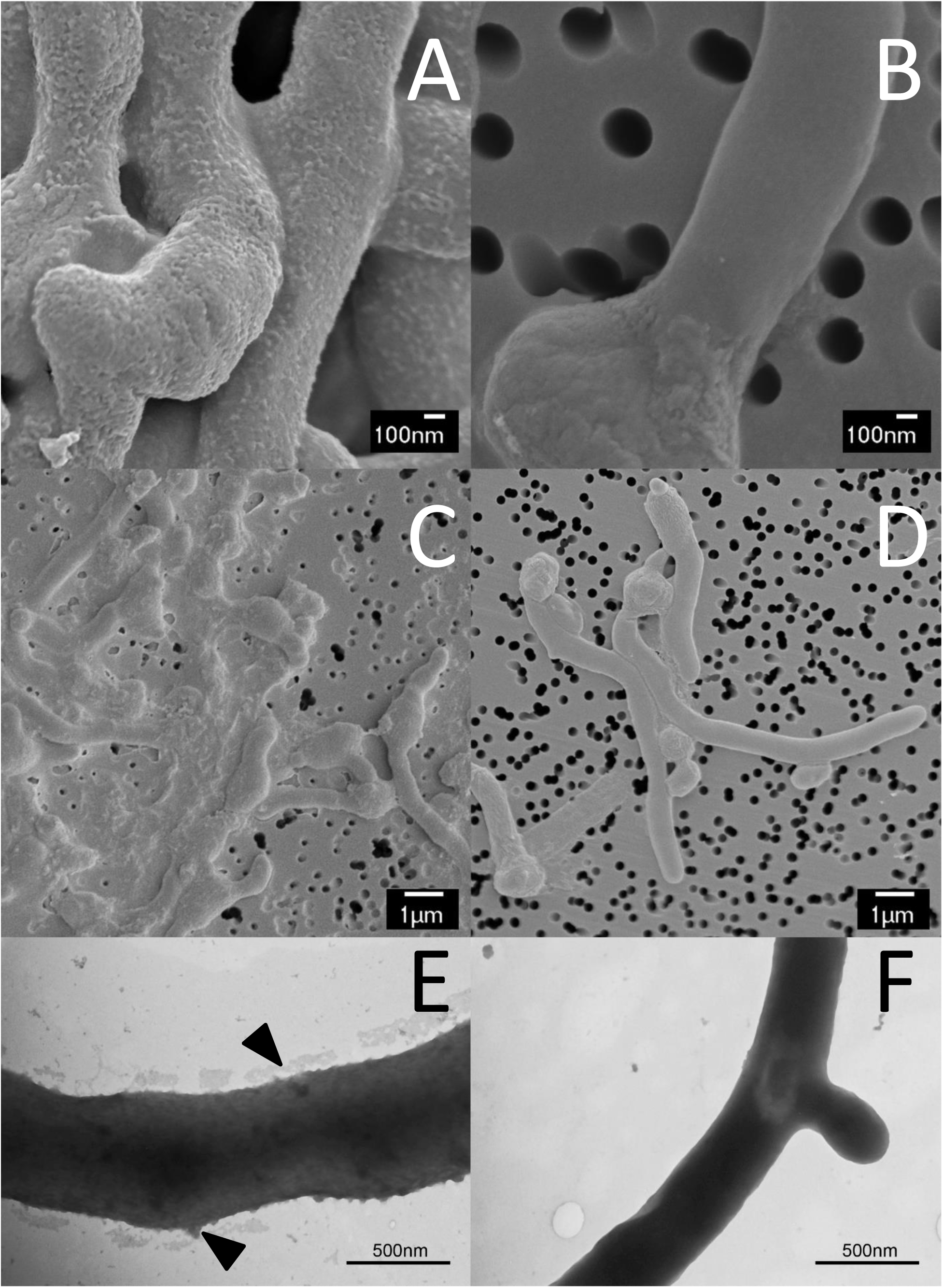
Demonstration of the extracellular layer produced by the *mat* genes. Cryo SEM of young vegetative mycelium (A-D) shows an abundance of extracellular material in wild type *S. lividans* covering the outside of hyphae (A) and between hyphae (C). This extracellular material is absent in the *mat* mutant (B and D). Negatively stained hyphae with tungsten acid, specific for polymeric substances, reveals a scabrous outside coating in wild-type hyphae (E) that is absent in the *mat* mutant (F). All strains were grown for 8 h in TSBS media in a shake flask.

### MatB correlates to the production of poly-N-acetylglucosamine

Bioinformatics analysis of MatA failed to identify known protein domains. MatB contains two functional domains, namely an intracellular glycosyltransferase type 2 (GT2) domain and a type 4 carbohydrate esterase (CE4) domain, connected by a predicted transmembrane helix. Sequences of glycosyltransferases and carbohydrate esterases were extracted from CAZy, which catalogs enzymes with characterized function, and assembled in a local database for Blast analysis. The glycosyltransferase domain of MatB returned PgaC from *E. coli* as the top hit (Table S1). *E. coli pgaC* encodes a glycosyltransferase that synthesizes Poly-β-1,6-*N-* acetylglucosamine (PNAG) (33). The next nearest homologs were enzymes with the same function in *Acinetobacter baumannii*, *Staphylococcus epidermis, Aggregatibacter actinomycetemcomitans and Actinobacillus pleuropneumoniae*, all with similar scores.

A similar blast comparison with the MatB carbohydrate esterase domain returned PgdA (BC_3618), a peptidoglycan N-acetylglucosamine deacetylase from *Bacillus cereus* as nearest characterized homologue (Table S1). Other top hits include a chitin deacetylase from *Caldanaerobacter subterraneus* and NodB proteins from *Rhizobium* species, all with similar scores. Interestingly, these enzymes all act on 1,4-linked oligo-chitin like substrates, in contrast to poly-β*-*1,6-N-acetylglucosamine glycosyltransferases. Submission of the MatB protein sequence to the PHYRE^2^ webserver, which models 3D protein structures using known crystal structures as inputs (34) returned a putative protein model where 86% of the residues could be modeled with more than 90% confidence (Figure 2A). The GT2 domain could be modeled with 100% confidence, using BcsA of *Rhodobacter sphaeroides* (PDB 4hg6) as template and the CE4 domain was modeled with 100% confidence, using a combination of six oligo-chitin/GlcNAc peptidoglycan deacetylases (PDB 2c1I, 1ny1, 4nz3, 1w17, 4m1B, 4l1G) (Figure 2B). The structure of the N- and C-termini and the transmembrane helix that connects the two domains could only be modeled with low confidence. Nearly all amino acid residues involved in binding of the UDP-sugar moiety in various PNAG biosynthetic glycosyltransferases are conserved in MatB (Figure 2C; Figure S2; Figure S3), while residues in the active site of the MatB CE4 domain had very high homology to those of oligo-chitin deacetylases (Figure 2D). Interestingly, neither PNAG deacetylases nor chitin synthases share a high homology with the respective MatB domains, and phylogenetic analysis using a maximum-likelihood tree build places MatB in the middle for both CE4 or GT2 domains (Figure S3). Taken together, bioinformatics analysis predicted that MatB synthesizes poly-*N*-acetylglucosamine, which could be either in the (1,4)- or in the (1,6)-configuration.

**Figure 2.**
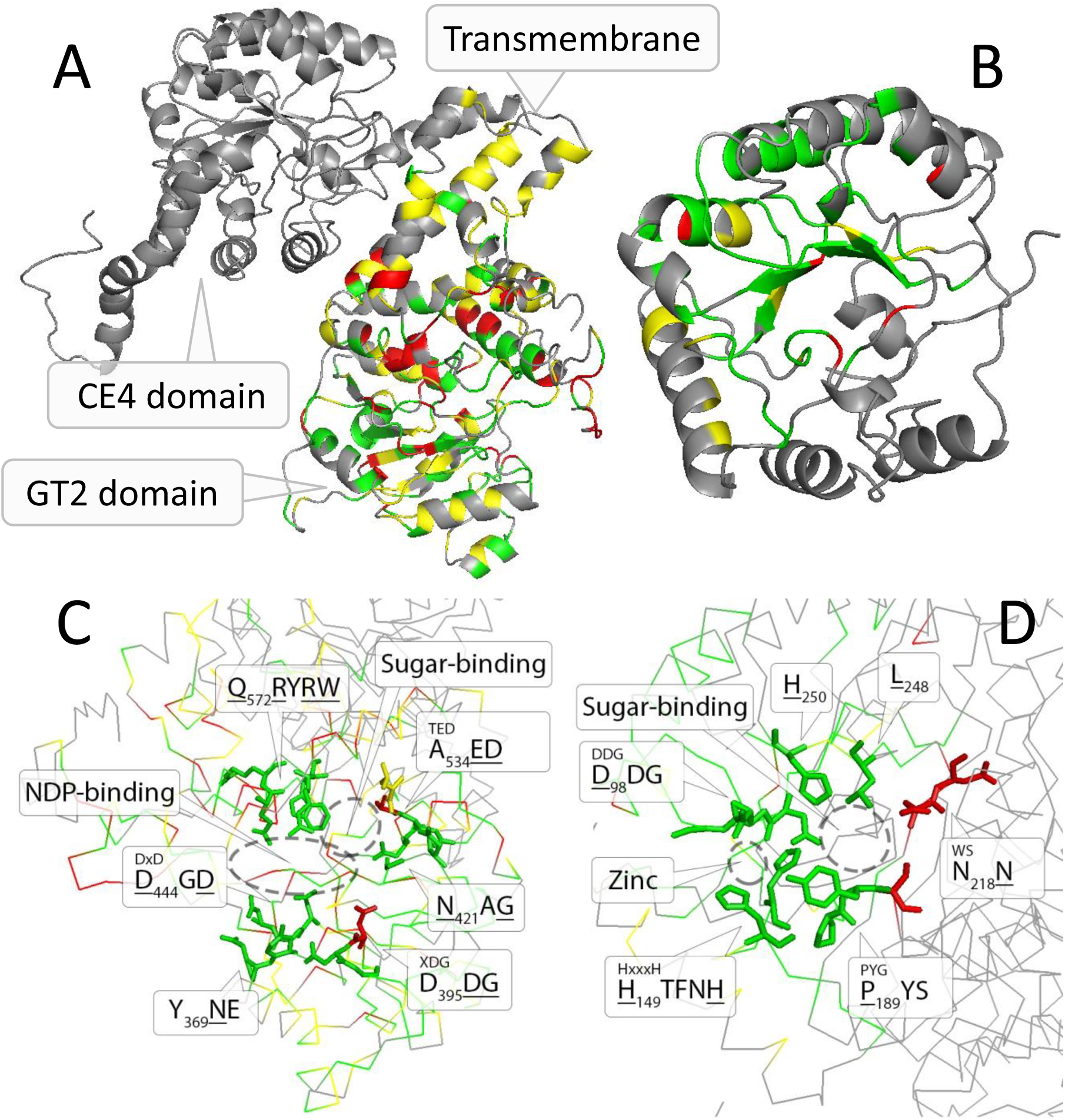
Structural model of the MatB protein. Model of predicted MatB using PHYRE^2^ where the two predicted domains were submitted as a whole and separately. The intracellular GT2 domain (aa 354-734) (A and C) was based on the cellulose synthase BcsA (4HG6), while the extracellular CE4 domain (aa 1-342) was based on a combination of 6 templates (PDB 2c1I, 1ny1, 4nz3, 1w17, 4m1B, 4l1G) (B and D). The coloring represents the conservation from the top 5 blast scores where green is 80% conserved in homologues and MatB, yellow is a conserved aa type and red represents 80% conservation in homologues, but different in MatB. Gray represents non conserved residues. The putative active sites and important amino acids involved in the enzymatic reactions are indicated in C and D for the glucosyltransferase domain and the carbohydrate esterase domain respectively.

### MatB produces a PNAG-like EPS

To analyze if the Mat proteins may be involved in the biosynthesis of (1,3-) or (1,4-) glycans, hyphae of *S. lividans* 66 and its *matB* null mutant were stained with calcofluor white (CFW) (35). Apical sites of both wild-type and *matB* mutant cells were stained with equal efficiency (Figure 3). This is contrary to the absence of staining in *cslA* null mutants, where the synthesis of (1,3)-or (1,4)-glycans is impaired (26, 29). CslA and its partner GlxA synthesize a cellulose-like polymeric substance, which is also involved in the aggregation of *Streptomyces* in liquid-grown cultures (36). This strongly suggests that MatAB do not synthesize (1,3)- or (1,4)-glycans.

**Figure 3.**
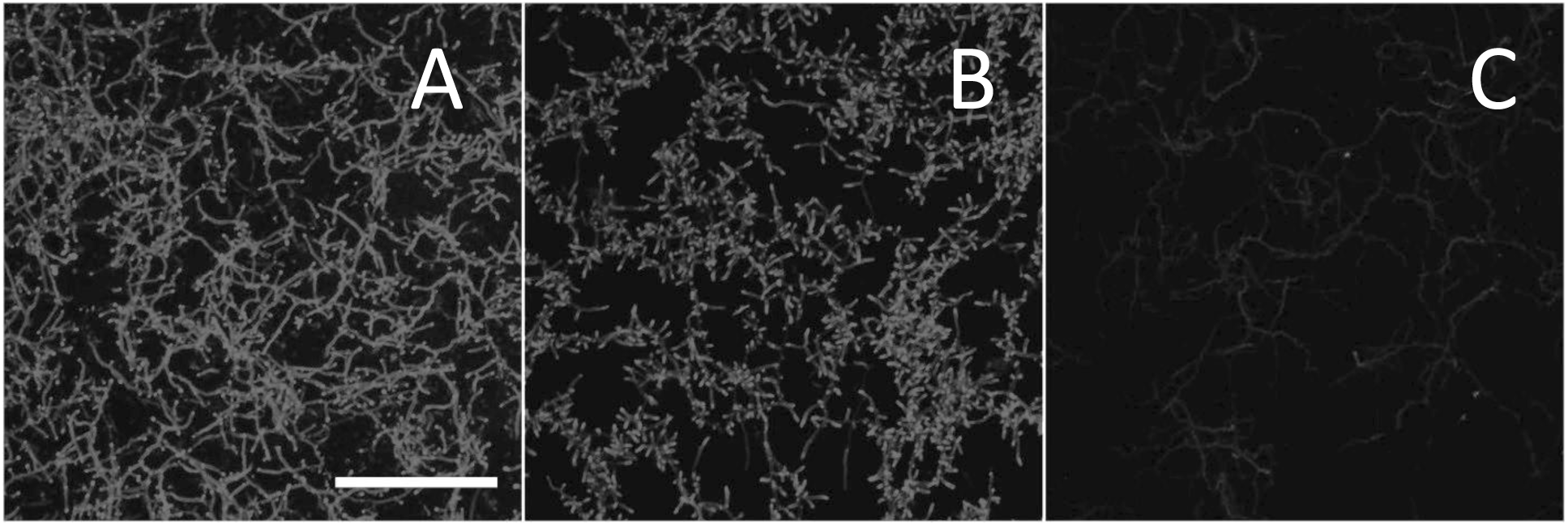
Calcofluor white staining. *S. lividans* (A), the *matAB* mutant (B) and Δ*cslA* (C) were stained with CFW to assess the presence of extracellular (1,3)- or (1,4)-glycans. The staining patterns indicate the presence of (1,3)- or (1,4)-glycans in both the parental strain and its *matAB* mutant, while it is absent in the *cslA* mutant. Scale bar equals 50 μM.

To further characterize the product of the MatAB enzymes, we used monoclonal antibodies (mAb F598) that specifically recognize both intact and deacetylated PNAG (37). Mycelia obtained from 6 h liquid-grown cultures of *S. lividans* 66 or its *matB* mutant were fixed in 4% PVA and incubated overnight with mAb F598. After washing and incubation with a fluorescently labeled secondary antibody conjugate, mounting fluid containing SYTO85 was added to stain the DNA and samples were then imaged with a fluorescence microscope (Figure 4). Wild-type cells were strongly stained with mAb F598, indicating the production of a PNAG-like polymer. Co-localization with the DNA stain SYTO85 suggests that most PNAG-like molecules are located on the cell surface. Conversely, immunofluorescence microscopy of *matB* null mutants with mAb F598 only resulted in background fluorescence, strongly suggesting that the Mat proteins indeed synthesize PNAG or a highly related PNAG-like polymer.

**Figure 4.**
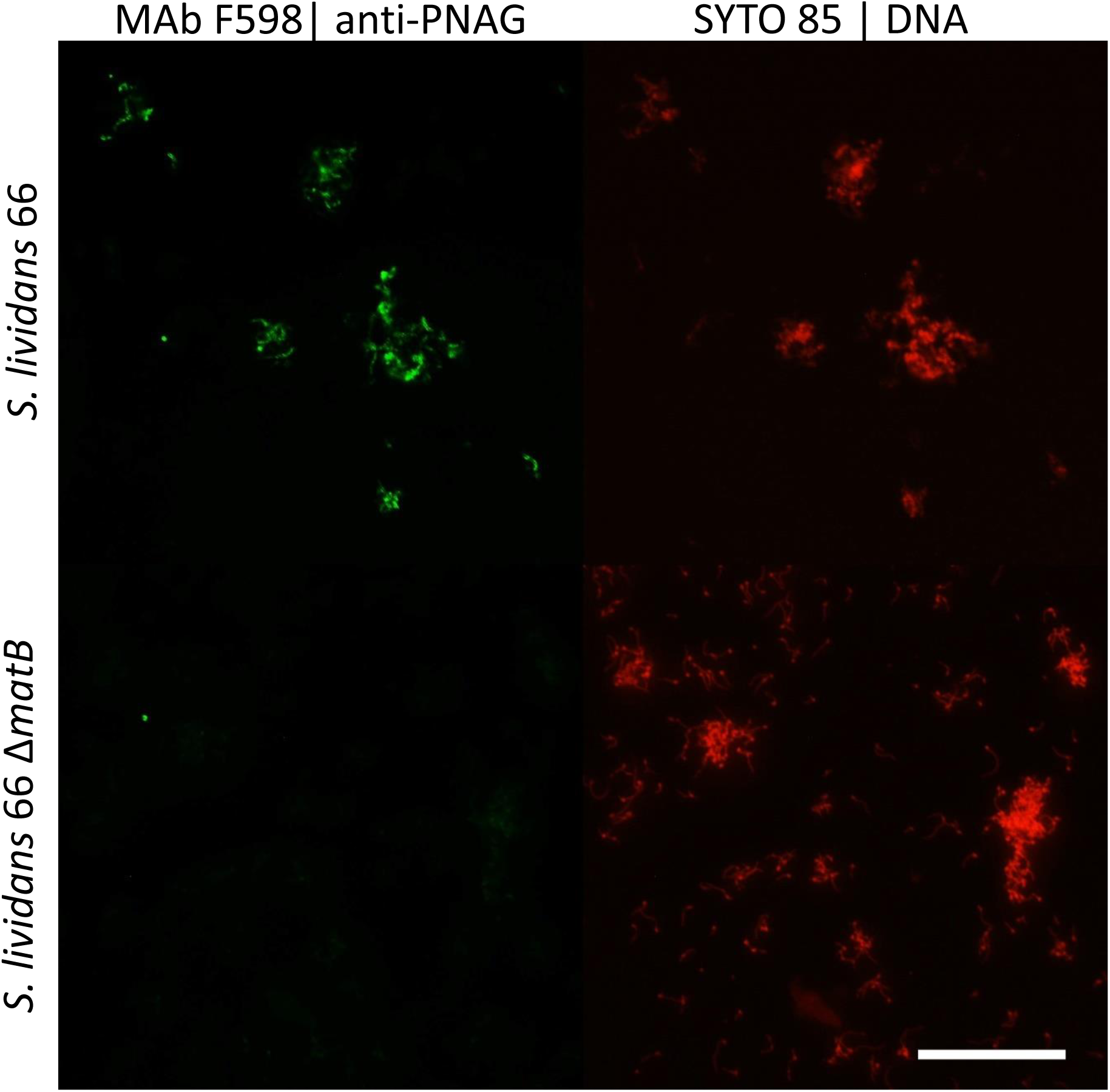
Immunofluorescence micrographs of *S. lividans* and its *matB* null mutant to identify extracellular PNAG. Young mycelia from 6 h old cultures of *S. lividans* 66 and its *matB* mutant were analyzed for the presence of PNAG with the specific monoclonal antibody mAb F598 and secondary anti-human IgG Alexa 488 conjugate. The presence of cells is indicated by the DNA binding dye SYTO 85. These experiment demonstrate that PNAG was produced by wild-type cells but not by *matB* mutants. Bar, 100 μm.

To further ascertain the presence of PNAG, the mycelia were treated for 2 h with a suspension containing either chitinases, cellulases or dispersin B. Only dispersin B, which specifically degrades PNAG (38), significantly affected EPS accumulation as visualized by SEM microscopy, further indicating that the extracellular *mat*-dependent EPS is indeed PNAG (Figure 5).

**Figure 5.**
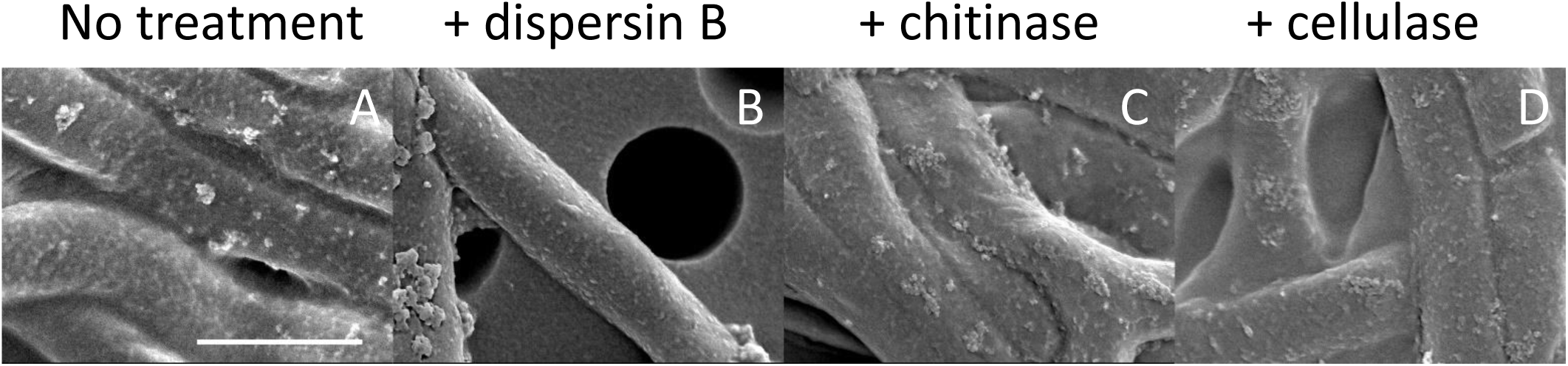
Effect of hydrolytic enzymes on the accumulation of EPS on hyphae of *S. lividans* 66. Mycelia of wild-type *S. lividans* 66 were treated with either 50 μg/ml dispersin B, 0.5 U/ml chitinase or 2 U/ml cellulase for 4 h. The biomass was imaged with cryoSEM to visualize the extracellular matrix. Note that treatment with dispersin B, which degrades PNAG, resulted in smooth hyphae. Bar, 1 μm.

### Mechanistic insight into matA and matB in relation to cslA and glxA

As mentioned above, the inhibition of mycelial pellet formation by *S. lividans* in liquid-grown cultures is not uniquely associated with *matA* or *matB*, but has also been observed when *cslA* and/or *glxA* are disrupted. When grown in liquid cultures the phenotypes of *cslA*, *glxA* or *matB* null mutants are phenotypically highly similar, with highly dispersed growth, highlighting the importance of both the *matAB* and *cslA-glxA* gene clusters for pellet formation. In an attempt to increase our understanding of how the two different EPSs might coordinate aggregation, we investigated the attachment behavior on hydrophobic and hydrophilic surfaces, via adherence assays on glass and polystyrene, respectively. Attachment of the *matA* and *matB* mutants to polystyrene attachment was similar to that of the parental strain, while attachment of *cslA* or *glxA* mutants was strongly reduced (Figure 6A). Conversely, attachment to glass is mostly depended on *matA* and *matB*, and was less affected in the *cslA* or *glxA* mutants (Figure 6B and Figure 7). This indicates that the EPS produced by CslA and GlxA plays a dominant role in adherence to hydrophobic surfaces, while the PNAG produced by MatAB is particularly relevant for adherence to hydrophilic surfaces.

**Figure 6.**
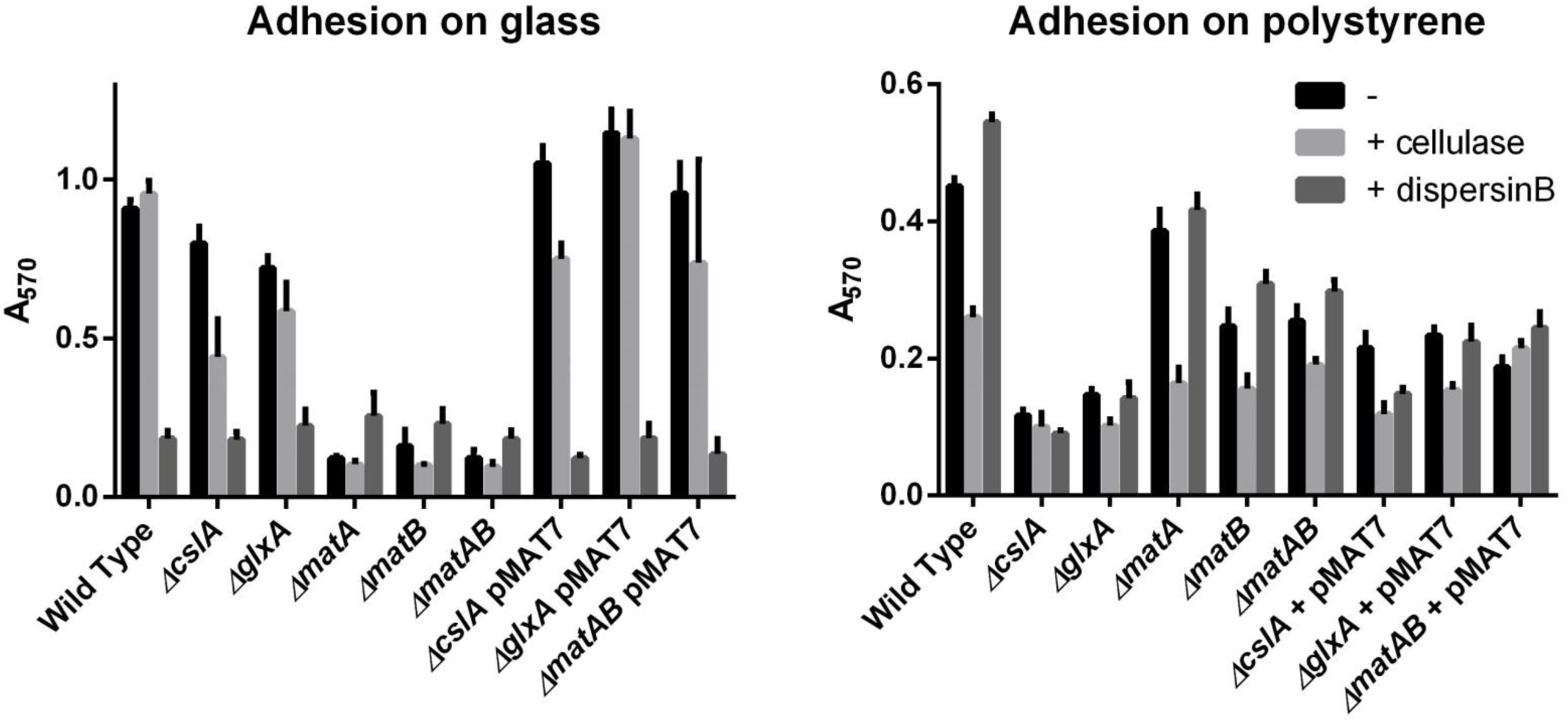
Quantification of attachment to solid surfaces. Surface attachment was quantified for *S. lividans* 66 and its respective c*slA*, *matA* or *matB* mutants with and without added Dispersin B or cellulase. Quantification was performed by staining attached cells with crystal violet and measuring dissolved crystal violet spectrophotometrically at 570 nm. The average and standard deviation of five independent wells are given. A) Surface attachment on glass from overnight growth B) Surface attachment to polystyrene after 7 days of growth.

**Figure 7.**
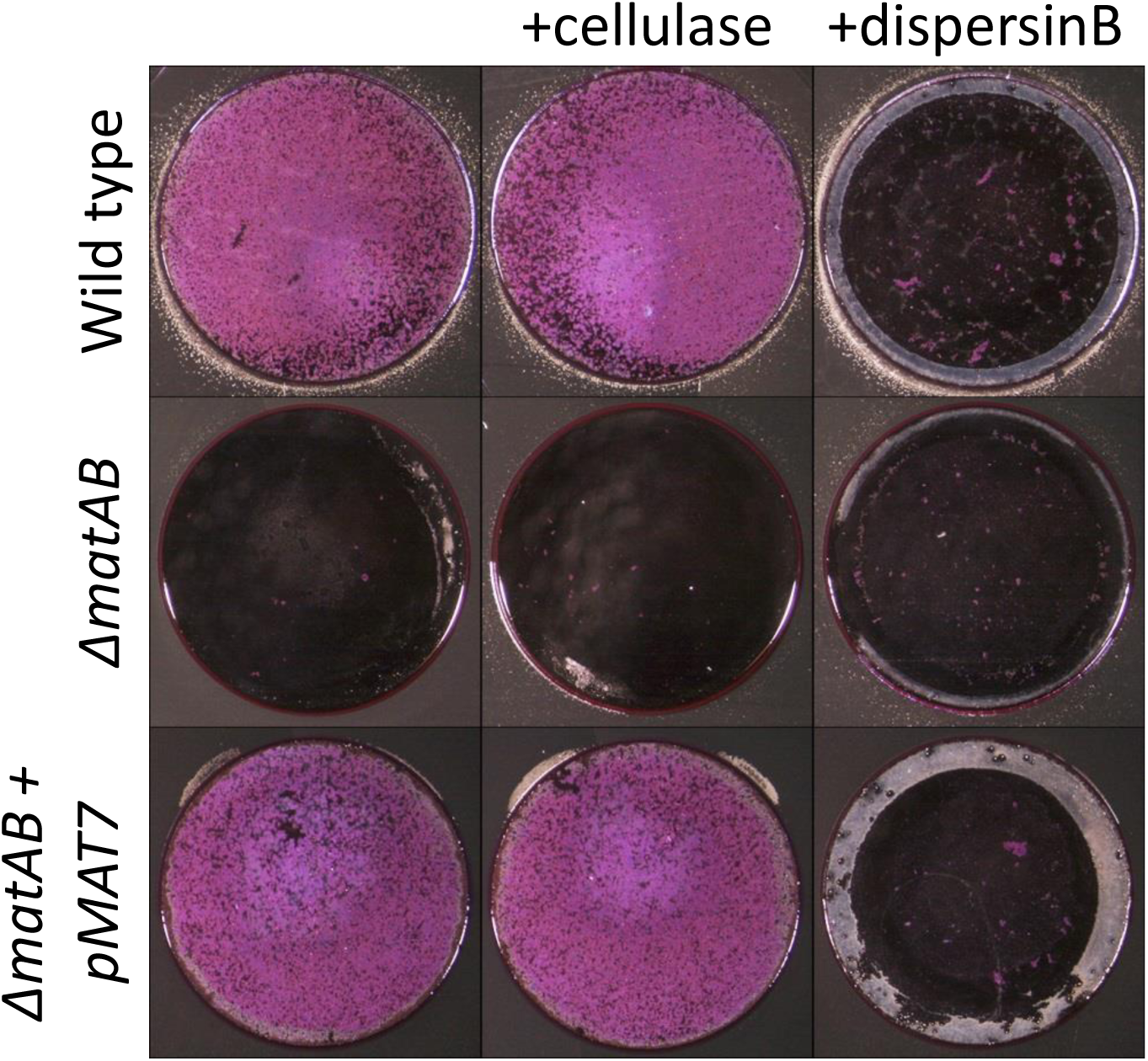
Visualization of adhesion to glass. *S. lividans* 66, its *matAB* null mutant and the *matAB* mutant complemented with pMAT7 grown for 20 h. Cellulase at a concentration of 0.2 U/ml had no effect on glass surface attachment, in contrast to 50 μg/ml dispersin B, which efficiently inhibited attachment. Complementation of the *matAB* mutant with pMAT7 restored attachment, which could in turn be antagonized again by the addition of dispersin B.

### MatAB expression is responsible for pellet formation

Pellet formation in shaken liquid cultures appears to depend on both hydrophilic and hydrophobic adhesive forces, as deletion of either *cslA* or *matB* prevented pellet formation (Figure 8 E and F). However, it might be more complicated than the sum of the two factors, as the addition of high concentrations of either cellulase (2 U/ml) or Dispersin B (100 μg/ml) was unable to alter the phenotypic characteristics of pellets (Figure 8 A-C). A mix of both enzymes did induce a morphological change, but did not prevent pellet formation, indicating that more factors determine the integrity of pellets than the *cslA*- and *matAB*-dependent EPSs (Figure 8 D). However, pellet formation could be restored to *cslA* mutants by the introduction of the pMAT7 construct, which over-expresses *matAB* from the strong constitutive *gapA* (SCO1947) promoter (Figure 8 G and H). Importantly, this MatAB-driven complementation of pellet formation by *cslA* mutants could be readily antagonized by the addition of dispersin B, which underlines that PNAG formation was responsible for the complementation. These data indicate that the enzymatic resistance of native pellets is the result of the complex composition of the extracellular matrix.

**Figure 8.**
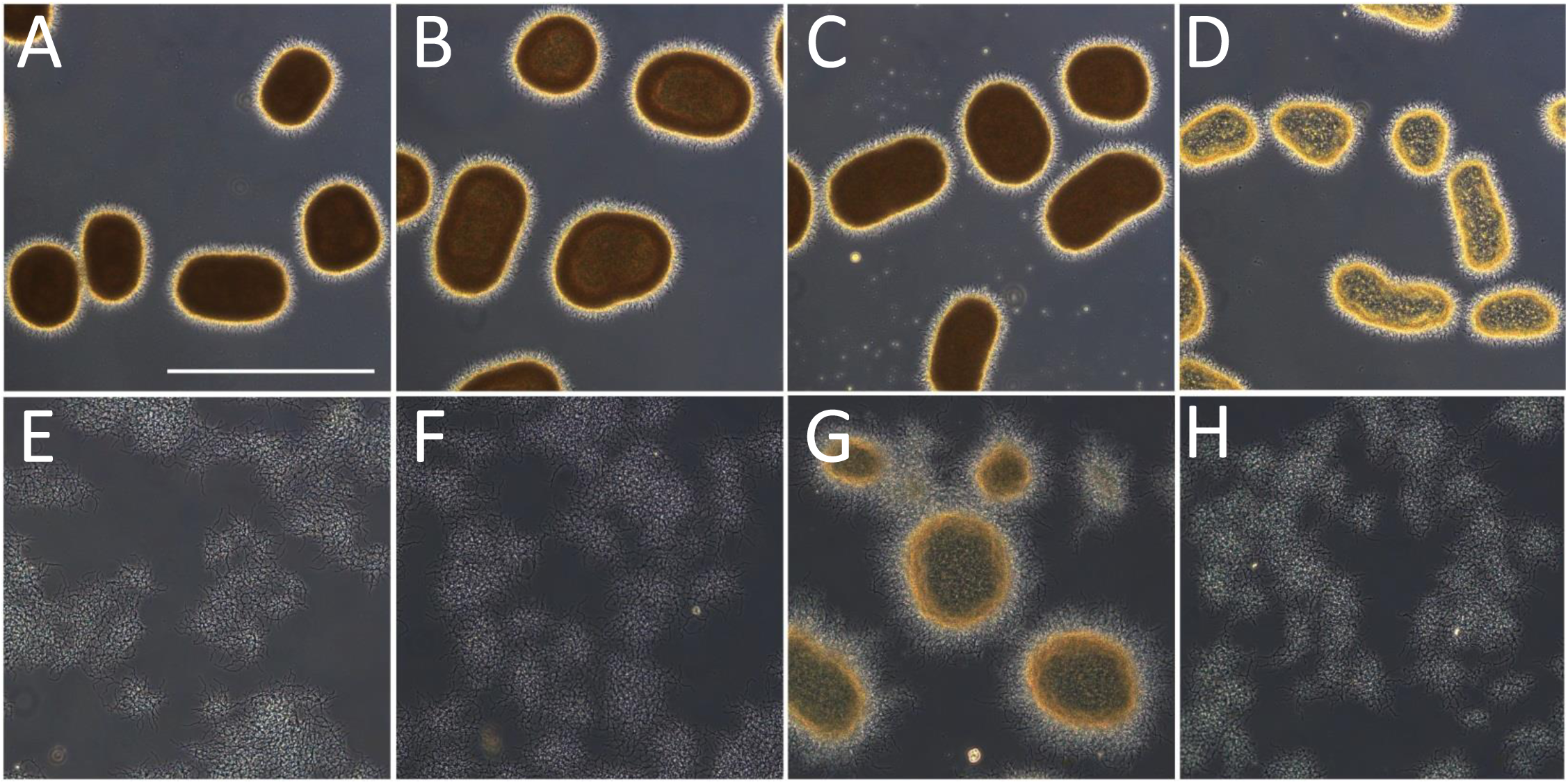
Effect of cellulase and dispersin B on mycelial morphology. Light micrographs show *S. lividans* 66 (A), *S. lividans* 66 treated with 2 U/ml cellulase (B), 100 μg/ml dispersin B (C) or both cellulase and dispersin B (D), the *matB* (E) and *cslA* mutants (F) and the *cslA* mutant harboring pMAT7 without (G) or with (H) added dispersin B. All strains were grown in TSBS medium for 24 h at 30°C. The effects on morphology were visualized by widefield microscopy. Bar, 500 μm.

Finally, we tested if addition of *matAB* to an actinomycete that does not have the genes on its chromosome would be sufficient to alter the mycelial morphology. As a test system we used the non-pelleting *Saccharopolyspora erythraea*. Introduction of plasmid pMAT7 into the strain indeed induced pellet formation, and again this phenotype was reversible by the addition of the PNAG-antagonizing enzyme dispersin B (Figure 9). The heterologous production of PNAG by MatAB shows that the presence of this polymer by itself suffices to induce pellet formation in filamentous actinomycetes, which opens new perspectives for morphological engineering approaches.

**Figure 9.**
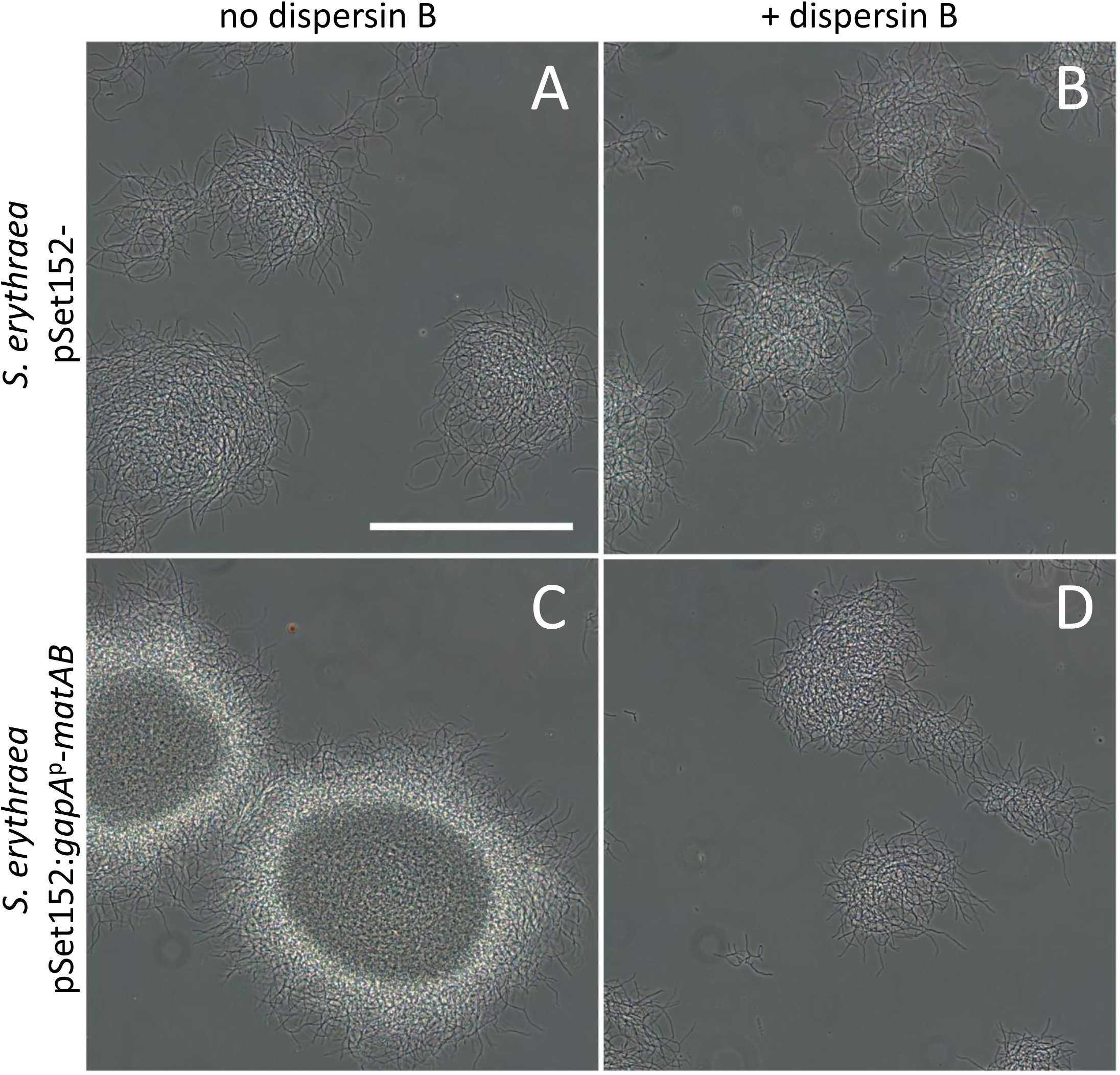
Effect of dispersin B on the mycelial morphology of *Saccharopolyspora erythraea*. *S. erythraea* transformants harboring the empty vector pSET152 (A and B; control) or pMAT7 (C and D) were grown for 24 h in TSBS media with (B and D) or without (A and C) 50 μg/ml dispersin B. The effects on mycelial morphology were visualized by widefield microscopy Note the increased aggregation of *matAB* transformants, which could be reversed by adding dispersin B. Bar, 200 μm.

## DISCUSSION

Members of the multicellular filamentous genus *Streptomyces* have an innate ability to self-aggregate in liquid-grown cultures, with the mycelia of several species forming dense pellets. This mode of aggregation contrasts with other surface-attached biofilms, which typically consist of aggregating single cells, held together by an extracellular matrix (4, 5). The matrix contributes to structural integrity of the multicellular community, while simultaneously providing protection against various stresses (Scher et al., 2005; Romero et al., 2010; DePas et al., 2013). Multiple matrix layers may be formed, with an outer layer containing a soluble EPS and a core with insoluble EPS and hydrophobic proteins (39). While matrices are usually mentioned in the context of biofilms, streptomycetes also make extracellular substances that contribute to morphology. Pellets of *S. coelicolor* were proposed to contain extracellular DNA (eDNA) and hyaluronic acid, and interference with these matrix components resulted in (partial) disintegration of mycelial pellets (Kim & Kim, 2004). In this work, we report the discovery of a similarly multi-layered and multi-component system in *Streptomyces*. Besides the previously identified cellulose-like EPS that is produced by the action of CslA and GlxA, we here show that the MatAB enzymes produce PNAG, which is also a well-known EPS that plays a role in biofilm formation by many planktonic bacteria (Wang *et al.*, 2004, Roux *et al.*, 2015, Mack *et al.*, 1996, Beenken *et al.*, 2004). These data strongly suggest that the formation of pellets by liquid-grown mycelia of streptomycetes may be based on the same principles as the formation of biofilms by planktonic bacteria. This apparently supports the hypothesis that hyphal growth of mycelial microorganisms may have evolved from the less permanent aggregation of single cells (4).

Adhesion assays with glass (hydrophilic) and polystyrene (hydrophobic) revealed that PNAG is primarily responsible for adhesion to hydrophilic surfaces (*i.e.* to glass), while the cellulose-like EPS promotes hydrophobic adhesion (to polystyrene). Both types of EPS play a crucial role in the maintenance of mycelial pellets in submerged cultures, and deletion of either *cslA* (29) or *matAB* (30) results in highly dispersed growth, although overexpression of *matAB* is sufficient for the formation of pellets by *cslA* null mutants. With both systems present, the obtained rigidity of the pellet architecture is more than the sum of its parts, as indicated by the resistance against the combination of dispersin B and cellulases. In *S. epidermis,* PNAG has been linked to adherence to both hydrophilic and hydrophobic surfaces (40). Why PNAG alone is not enough for hydrophobic adherence in *Streptomyces* requires further investigation, but might be related to the larger multicellular size of the organism, requiring a greater force for adherence. Although not understood in detail, hydrophobic adherence of *Streptomyces* is likely the result of a more complicated system, which besides the product of CslA also involves the hydrophobic chaplin proteins (27). As the chaplins are expressed late in the life cycle, involvement of these proteins might also explain why strong attachment to polystyrene is a multi-day process (27). Chaplins are also involved in pellet formation, although the absence of the chaplin layer has less severe morphological consequences than lack of the cellulose-like or PNAG-based EPSs (unpublished data). The chaplin based hydrophobic forces, likely located in the core of a pellet, might contribute to strengthening the pellet, but cannot fully explain the role of CslA/GlxA in cellular aggregation. Earlier work indicated that the polymer produced by CslA/GlxA plays a role in stabilization of the tip complex (29), which might explain its pleiotropic involvement in multiple systems throughout the life cycle. We can speculate that aggregation by PNAG is to some extent dependent on the proper organization of the tip complex supported the polymer produced by CslA/GlxA. It is our hope that high resolution spatial co-localization studies of CslA/GlxA, the chaplins and MatAB in native pellets, currently in progress in our laboratory, shed light on the involvement of these systems and their interactions in controlling the mycelial architecture.

Understanding how hyphal aggregation and pellet formation is controlled brings us one step closer to controlling the morphology of streptomycetes in liquid-grown cultures, which is highly relevant for tuning the morphology to product formation (17, 41) After all, several antibiotics such as erythromycin (produced by *Saccharopolyspora erythraea*) and actinorhodin (by *S. coelicolor*) are solely produced when a minimum pellet size is achieved, while enzyme production is typically favored by fast growing and fragmenting hyphae (42, 43). Previously, primarily genetic approaches were followed to tune mycelial morphology. Over-expression of *ssgA,* which controls hyphal morphogenesis and activates cell division (20, 21), effects fragmentation of the hyphae by enhancing cell division, resulting in increased growth and enzyme production rates (42). However, a drawback to this approach is the major effect of SsgA on the cell cycle, with enhanced sensitivity to shear stress as a result. In this respect morphological engineering targeting extracellular glue-like substances such as PNAG- and cellulose-like EPSs, offers an attractive alternative, as the effects on the internal physiology are likely minimal. Thus, besides their high relevance for our ecological understanding of how streptomycetes grow and attach to surfaces in their natural environment, the insights gained by this work may also help to develop novel technologies that improve growth and productivity of streptomycetes.

## MATERIALS AND METHODS

### Bacterial strains and plasmids

The bacterial strains and plasmids used in this study are listed in Table S2. *E. coli* JM109 (44) was used as a routine host for plasmid construction. The native *matAB* locus and *gapA* promoter region were PCR-amplified from the *S. coelicolor* genome as described using primers SCO2963_F, SCO2962_R and PSCO1947_F, PSCO1947_R respectively (Table S1). The *matAB* locus was cloned as an EcoRI/BamHI fragment into the integrative vector pSET152 (45) and the promoter region was placed in front of the *matAB* locus as an EcoRI/NdeI fragment, resulting in construct pMAT7. Conjugative plasmid transfer to *Streptomyces* was done using *E. coli* ET12567 (46) harboring pUZ8002 as the host (47).

### Culture conditions

Streptomycetes were grown in shake flasks with a coiled stainless steel spring in 30 ml tryptic soy broth (Difco) with 10% sucrose (TSBS). Cultures were inoculated with 10^6^ cfu/ml and grown at 30°C. To assess growth in the presence of hydrolytic enzymes, strains were grown in 96-well plates where the agitation was facilitated by a Microplate Genie Digital mixer (Scientific Industries, USA) set to 1400 rpm, which was found to reproduce native morphologies at a micro scale (DVD and GVW, unpublished data). Dispersin B (100 μg/ml), cellulase (SigmaAldrich, C1184) (2 U/ml) or chitinase (SigmaAldrich C8241) (0.5 U/ml) were added during growth to degrade EPS. The strains were observed after 24 h of growth by wide field microscopy.

### Bioinformatics

The genomes of *Streptomyces coelicolor* A3(2) M145 (48) and *S. lividans* 66 (49) have been published. Protein domains were annotated using the conserved domain search v3.14 (50), using default settings. Homology searches were performed using the local Blast+ software v2.2.30 (51). A BlastP database was built from the amino acid sequences of all characterized type 2 glycosyltransferases and type 4 carbohydrate esterases listed in the CAZy database (www.CAZy.org). The amino acid sequences were retrieved from the Uniprot database (www.uniprot.org). *In silico* structure prediction of MatB was performed with the Protein Homology/analogy Recognition Engine Version 2 (PHYRE^2^) (34). Structural analysis and alignment was performed in Pymol (v1.7.4). Sequence alignments were done in MEGA (v7.0.9) using the ClustalW algorithm (52). Maximum likelihood trees were constructed using default settings and a bootstrap with 500 iterations.

### Production and isolation of dispersin B

Dispersin B from *Aggregatibacter actinomycetemcomitans* ATCC 29522 was produced and purified as described (38). The specific activity, determined as the amount of enzyme needed to hydrolyze 1 μmol 4-nitrophenyl-β-D-N-acetylglucosaminide per minute in 50 mM sodium phosphate buffer (pH 5.5) 100 mM NaCl was 570 U /mg protein.

### Calcofluor white staining

The presence of (1-3) or (1-4) glycans was assessed by calcofluor white staining (53). Strains were grown over night in 8-well microscope chambers (LabTech II) in 300 μl TSBS medium. 30 μl calcofluor white (CFW) solution (Sigma Aldrich) was added and after 5 min incubation the samples were imaged on a Zeiss LSM5 Exciter/ Axio observer with a 405 nm laser, a 405/488 nm beam splitter and 420-480 nm bandpass filter (54).

### Immunofluorescence

Immunofluorescence microscopy was performed as described (10), with small adaptations. In short, *S. lividans* 66 was grown in TSBS media for 6 hours at 30⁰C. A 50 μl culture aliquot was spotted inside circles drawn with a PAP pen on adhesive microscope slides (Klinipath, The Netherlands). After 15 min the media was removed gently and the cell layer was air dried for 10 min and fixed with 4% paraformaldehyde in PBS for 15 min. After washing samples twice with PBS, monoclonal antibodies against PNAG (mAb F598) were added to a final concentration of 10 μg/ml in PBS with 0.1% BSA-c (Aurion, the Netherlands) and samples incubated for 16 h at 4⁰C. The samples were then washed three times with PBS with 0.1% BSA-c and fluorescently-labeled goat-anti-human IgGs (Life Technologies) added to a final concentration of 4 μg/ml and incubated in the dark for 2 h. After washing twice with PBS with 0.1% BSA-c, some PBS with propidium iodide at a concentration 1 μg/ml was added and the samples were imaged on an axiovision Zeiss microscope equipped with a mercury lamp.

### Cryo scanning electron microscopy

Mycelia from cultures grown for 6 h, fixed by 1,5% glutaraldehyde and immobilized on isopore membrane 0.8 μm filter discs (Millipore) by pushing the liquid through using a syringe and placing the filter in a filter holder. The discs were cut to size and placed on the SEM target immobilized with Tissue Tek® and quickly frozen in liquid nitrogen slush and transferred directly to the cryo-transfer attachment of the scanning electron microscope. After 10 minutes sublimation at -90 ˚C specimens were sputter-coated with a layer of 2 nm Platinum and examined at -120 ˚C in the JEOL JSM6700F scanning electron microscope at 3 kV as described (55).

### Negative stain TEM microscopy

For negative staining, 5 μl of young mycelium was placed on a copper TEM grid and air dried for 15 min. The cellular material was stained with 3% PTA for 5 min, followed by 5 times washing with miliQ. The samples were placed in a JEOL 1010 transmission electron microscope and observed at 60 kV as described (56).

### Adhesion Assays

Attachment of strains to polystyrene surfaces was tested as described (57). In short 10^6^ CFUs/ml were inoculated into 4 ml NMMP (47) without polyethylene glycol and casamino acids, using 2% mannitol as the sole carbon source. After 5 days at 30°C the standing cultures were stained with crystal violet. After washing the attached cells were quantified by extracting the crystal violet with 10% SDS and measuring the absorption at 570 nm. Attachment to glass surfaces was tested in a similar fashion, using glass bottom 96 wells plates (Greiner Bio-One, Austria) and 200 μl NMMP medium without polyethylene glycol, but with 0,5% casamino acids and 2% glucose as the carbon source. These were cultivated overnight at 30°C and the attached biomass was quantified as for polystyrene.

## ACKNOWLEDGEMENTS

We like to thank Marleen Janus for the supply of *A. actinomycetemcomitans* gDNA and Ellen de Waal for the production of dispersin B. The research was supported by VIDI grant 12957 and VICI grant 10379 from the Netherlands Technology Foundation STW to DC and GVW, respectively.

